# Genome ARTIST_v2 software – a support for annotation of class II natural transposons in new sequenced genomes

**DOI:** 10.1101/2020.10.30.360610

**Authors:** Alexandru Al. Ecovoiu, Iulian Cristian Ghita, David Ioan Mihail Chifiriuc, Iulian Constantin Ghionoiu, Andrei Mihai Ciuca, Alexandru Marian Bologa, Attila Cristian Ratiu

**Author notes:** Corresponding author: Alexandru Al. Ecovoiu.

## Abstract

Transposon annotation is a very dynamic field of genomics and various tools assigned to support this bioinformatics endeavor were reported. Genome ARTIST (GA) software was initially developed for mapping artificial transposons mobilized during insertional mutagenesis projects. Now, the new functions of GA_v2 qualify it as an effective companion for mapping and annotation of class II natural transposons in assembled genomes, contigs or sequencing reads.

Tabular export of mapping and annotation data for subsequent high-throughput data analysis, the export of a list of flanking sequences around either the coordinates of insertion or around the target site duplications (TSDs) and generation of a consensus sequence for the respective flanking sequences are all key assets of GA_v2.

Additionally, we developed two accompanying short scripts that enable the user to annotate transposons existent in assembled genomes and to use various annotation offered by FlyBase for *Drosophila melanogaster* genome.

Herein, we present the applicability of GA_v2 for a preliminary annotation of the class II transposon P-element in the genome of *D. melanogaster* strain Horezu, Romania, which was sequenced with Nanopore technology in our laboratory. Our results point that GA_v2 is a reliable tool to be integrated in pipelines designed to perform transposon annotation in new sequenced genomes.

GA_v2 is open source software compatible with Ubuntu, Mac OS and Windows and is available at https://github.com/genomeartist/genomeartist and at www.genomeartist.ro.

## Introduction

Over the last several years, the bioinformatics tools that are able to detect and map transposable elements (TEs) insertions operate primarily on high-throughput sequencing data (Bergman and Quesneville, 2007; Vendrell-Mir *et al*., 2019).

Initially designed for mapping artificial transposon insertions, Genome ARTIST v1.19 (GA_1.19) differs from most tools dedicated to transposable elements analysis due to its intuitive and clear graphical interface, as well as its high mapping efficiency and accuracy (Ecovoiu *et al*., 2016). Nowadays, massive efforts are made to sequence the genomes of many different organisms that are subsequently stored in dedicated databases, hence providing huge amounts of data to be analyzed. Advances in TEs annotation projects require new and improved bioinformatics approaches in order to develop more specific and precise tools for best suiting the annotators’ needs.

Considering the present scientific context, we improved Genome ARTIST, in order to make it able to cope with the new trends in TEs research involving high-throughput approaches and automating data exploration.

Here, we present Genome ARTIST v2 (GA_v2), a version that contains several additional features which proved to be very effective when addressing high-throughput data analysis projects. The new implemented functions allow users to retrieve subsequences (TSD and flanking sequences) from specific genomic coordinates without prior alignment of the query sequences and to extract TSD and flanking sequences in FASTA format consecutive to the alignment of a set of transposon-genome junction sequences versus reference sequences. GA_v2 is also able to export a data table that includes the insertions coordinates, hit genes, upstream/downstream genes and the TSD sequence generated by each insertion.

## Methods

Implementation of the new functions in GA_v2 (Genome ARTIST v2.0.2 - DOI: 10.5281/zenodo.4139251) has been performed in the *Java* programming language. Using *Apache Ant* compiler and OpenJDK 8, GA_v2 code can be compiled on Linux, Mac OS and Windows.

For GA_v2 installation and run the detailed instructions are available at Ecovoiu *et al*. (2016), in the corresponding Manual and at website (http://www.genomeartist.ro/).

## Results

### I. The new features of GA_v2

#### 1. GA_v2 upgrades relative to computer usage

**1.1** An important upgrade from GA_1.19 to GA_v2 is the cross-platform compatibility. Our tests revealed that the new version is completely functional on Ubuntu/Mint, Mac OS and Windows operating systems, allowing the users to run the tool on their favorite OS, without the employment of virtual machines.

**1.2** GA_v2 evolved to a 64-bit one as compared to GA_v1.19, a feature that allows the user to harness more than 4 GB of RAM. This upgrade is a key one since manipulation and analysis of medium to large size genomes usually requires computers with at least 32 GB of RAM. In our hands, GA_v2 permits a smoother aligning/mapping experience, noticeable mainly when big genomes, such as the mammalian ones, are used as reference sequences.

#### 2. GA_v2 upgrades relative to the export of annotation data and sequence manipulations

**2.1** All of the annotations associated with the mapping results and depicted in the graphical interface are now exportable in a tabular form, which permits various subsequent manipulations of the mapping/alignment data. Thus, in addition to the insertion coordinates and chromosomal localization, GA_v2 also exports the name of the associated transposon loaded to Transposon files, the hit gene and the upstream/downstream genes in the vicinity of the insertion, the alignment score and target site duplication (TSD) nucleotide sequences of each TE insertion. By loading GA_v2 with a reference genome annotated with both genes and natural transposable elements, insertions associated with other transposable elements or any other annotated feature can also be identified (Additional figure 1).

**Figure 1.**
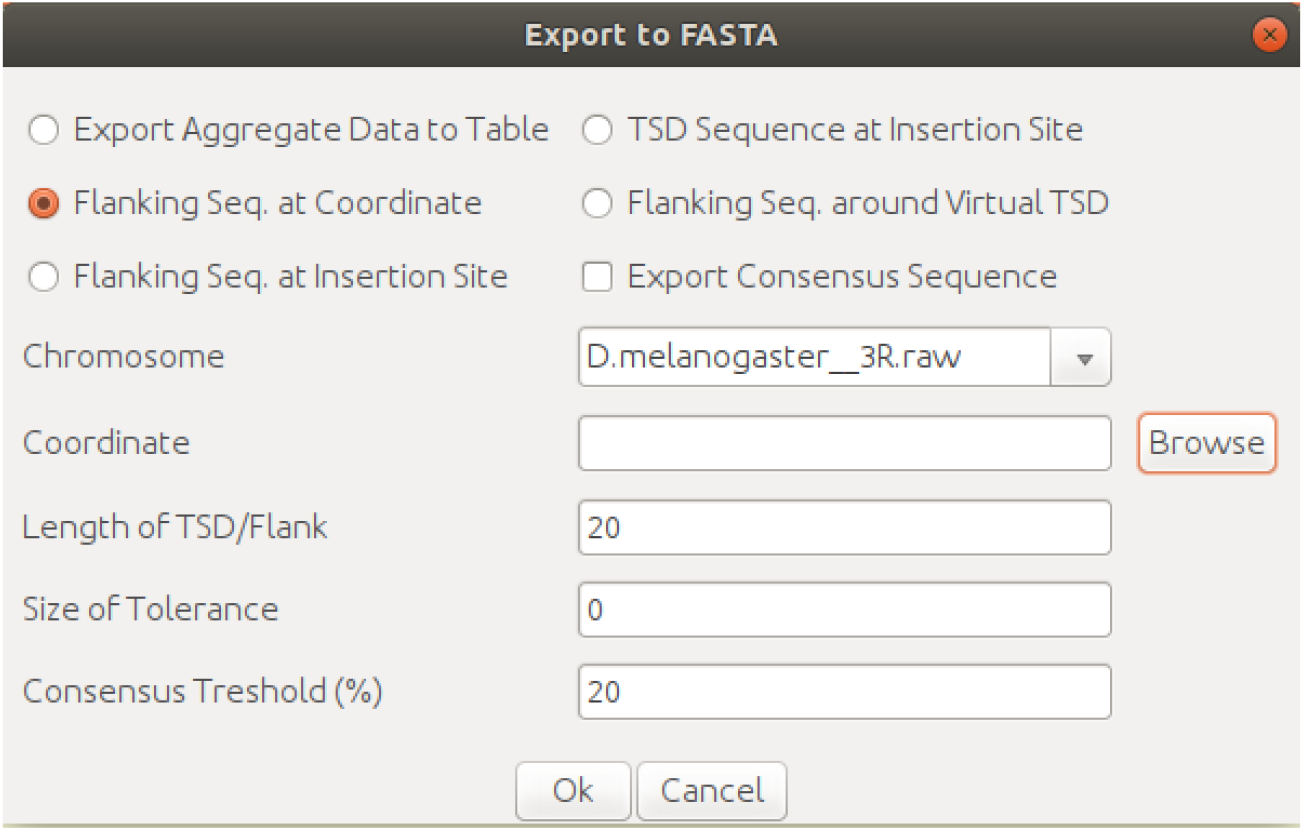
The *Flanking seq. at coordinate* function that enables the export of specified length nucleotide sequences centered on coordinates provided either directly, one by one, or by uploading a predefined list through the *Browse* option.

**Figure 2.**
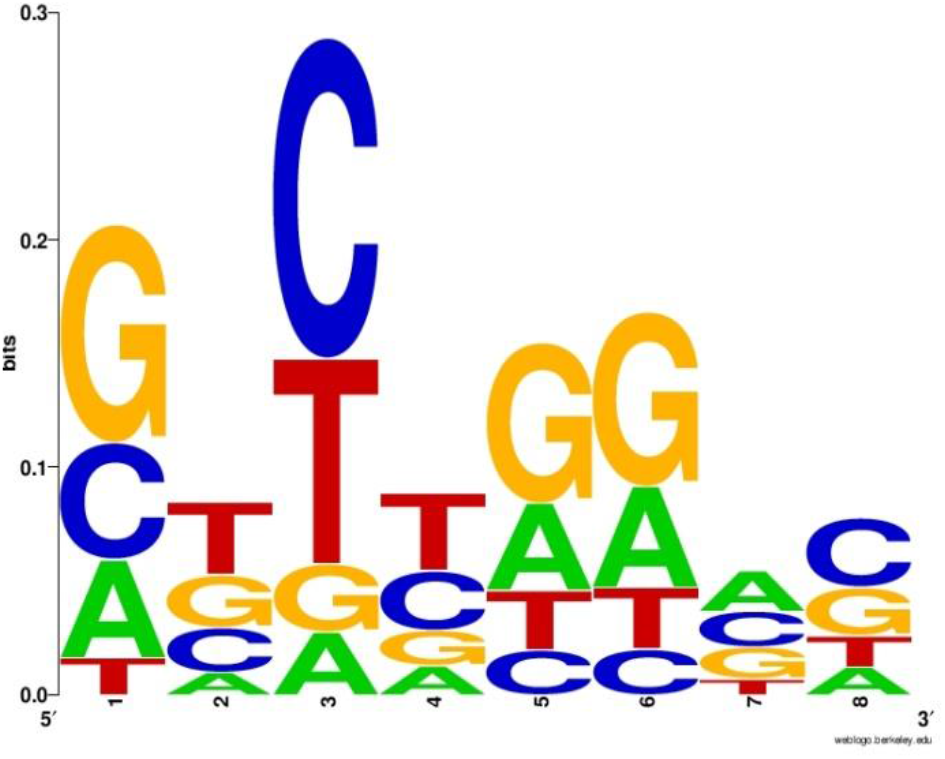
The sequence logo generated by WebLogo using the .fasta file containing a set of 125 TSD sequences from the Horezu local *Drosophila melanogaster* strain.

**2.2** GA_v2 is able to retrieve symmetric sequences of customizable lengths, which flank any number of input target nucleotide coordinates of a reference sequence. The flanking sequences are exported as .fasta files, so they may be readily used for various subsequent analyses, such as multiple alignments and consensus motifs detection. For this purpose, the user is expected to press *Export to FASTA* button.

Flanking sequences can be obtained around border nucleotides identified by the algorithm or around user specified coordinates. For the later, the *Flanking Seq. at Coordinate* option must be selected and then a chromosome must also be chosen from the drop-down list (Figure 1). A single coordinate can be specified in the adequate field or a .txt file can be loaded, where each line contains a coordinate.

A value for the length of the flanking sequence can be set in the *Length of TSD/Flank* field, where the default value of this parameter is 8, standing for the TSD length generated by insertion of the P-element in *D. melanogaster* genome. For example, if one changes this value to 20, then two local subsequences of 20 nucleotides centered on each coordinate of interest are considered for constituting the output sequences (one of the subsequences contains the respective query coordinate). Therefore, a list of specific contiguous sequences of 40 nucleotides will be extracted and exported as a .fasta file (the output file) for each of the nucleotide coordinates present in the input file.

If the exported flanking sequences are relatively short (a few tens of nucleotides) and are to be further aligned with GA_v2 for various purposes, such as local homology and oligonucleotide specificity, we recommend to set the value of *picking depth* parameter at a value around 100 (the default value is 1). Otherwise, alignments standing for other coordinates in the reference genome may be reported in the list of results. Obviously, if there is only one reference chromosome sequence existent in the database, this precaution is not necessary.

Similarly, consecutive to a post-alignment analysis either a list of TSD nucleotide sequences or a list of TSDs encompassed by flanking sequences of preset lengths may also be exported as a .fasta file. In this case, the input contains a list of junction sequences instead of chromosomal coordinates, each containing at least a transposon-genome or a transposon-transposon border. As mentioned above, the length of a TSD is customizable.

Using the *TSD Sequence at Insertion Site* function on data collected from the genome of a local *Drosophila melanogaster* strain from Horezu, Romania (BioProject/NCBI, accession PRJNA629549), we generated a list of 125 P-element 8-bp TSD sequences that were further loaded in WebLogo (Crooks *et al*., 2004). Thus we obtained a TSD consensus highly similar to the ones previously computed by Liao *et al.* (2000) and by Linheiro and Bergman (2012).

**2.3** Starting from a list of TSDs or flanking sequences, a consensus sequence can be generated by GA_v2 using IUPAC nucleotide ambiguity code. The user must load a list of junction sequences in GA_v2, then activate *TSD Sequence at Insertion Site*, check *Export Consensus Sequence* and press *Ok* button. The consensus sequence is exported in FASTA format and may then be used to scan against specific databases to find related consensus/motifs counterparts.

The *Consensus Threshold* value is set by default at 20% and is customizable. This value means that a specific nucleotide should be present in the same position in at least 20% of the exported sequences in order to be considered when selecting an IUPAC symbol for the consensus sequence. Obviously, if the value is set at 100%, the TSDs or the flanking sequences must be identical in order to obtain a consensus sequence out of them. But in this case, the consensus sequence would rather be a strict repetitive motif sequence, as it is the case of an invariable restriction site.

For example, the list of 125 P-element 8-bp TSD sequences particular to Horezu strain were analyzed by GA_v2 and the consensus sequence **VBYYRRVS** was generated.

#### 3. GA_v2 upgrades relative data visualization

**3.1** In GA_v2, the coordinate of insertion at the junction border (terminal genomic nucleotide or TGN) is highlighted in bright green, helping the user to identify the proper coordinate of insertion, mainly when the iPCR-derived sequence has transposon - genome - transposon or transposon - transposon - transposon (the particular case of self-insertions) structures.

**3.2** GA_v2 saves mapping/aligning results in Results folder with the date and a name chosen by the user, not with aleatory names as in Genome ARTIST v1.19. The results may be reviewed even with an empty GA_v2 package (containing no reference sequences). In addition, multiple results may be selected to be opened in the graphical interface.

**3.3** GA_v2 package contains two new scripts, which are available in the folder *scripts*. In contrast to GA_v1.19, the *FlyBase_annotation.sh* script that is found in the folder FlyBase allows the user to simultaneously associate multiple sets of annotations with the reference sequence of *D. melanogaster* genome. It uses .fasta.gz files downloaded from FlyBase that contain annotations of *D. melanogaster* genome such as genes, transposons, pseudogenes, predicted features, three prime UTRs and several others. The script simply needs to be run in the folder where are located the .fasta.gz files that contain the annotations and it will generate a _gene.fasta file corresponding to each chromosomal arm. Generated FASTA files are loaded in GA_v2 together with the .raw files (previously downloaded from FlyBase FTP server).

Therefore, the location of a transposon insertion may be interpreted in a more complex context of the genomic local landscape revealed by graphical interface of GA_v2. For example, insertion of a transposon into a different transposon located in its turn close to or overlapping a gene is easy to be noticed in GA_v2 (Figure 3). As presumed, this feature in functional only for annotation data available in FlyBase (Thurmond *et al.,* 2019).

**Figure 3.**
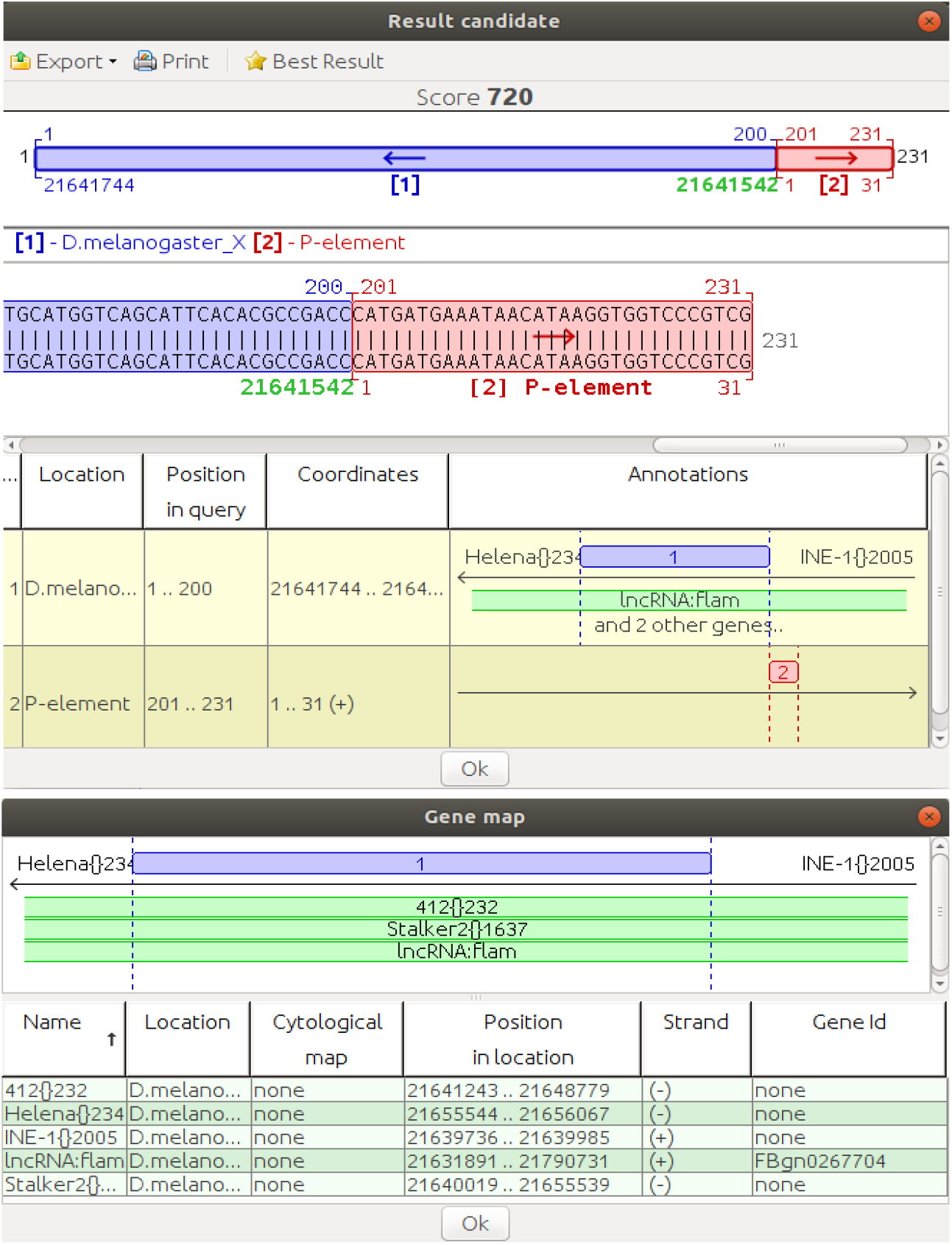
Partial alignment of Nanopore sequencing read (*D. melanogaster* strain from Horezu, Romania). The insertion is located in a non-protein coding gene that overlaps two natural transposable elements, 412{}234 and Stalker2{1637} (according to *D. melanogaster* r6.36 reference genome). In the lower panel, two other natural transposons are depicted as located upstream and downstream of the highlighted genomic region.

The second script, *TE_annotator.sh*, automatizes manual annotations of transposons, allowing the user to define IRs, genes and other annotations within the mobile element. The script must be run from “root folder”/resources/gene/ and can be run in the Ubuntu terminal. It is important for the name of the transposon indicated in the appropriate terminal window to coincide with the name used in GA_v2 (we assume that one or more transposons have been preloaded in GA_v2 database). Before allowing the user to annotate the desired features, the script requires the coordinate intervals/ranges of the 5’ and 3’ ends to be specified. Figure 4 shows the terminal window into which the annotation of the natural transposon P-element were implemented using the *TE_annotator.sh* script, while in Figure 5 the graphical implementation of these annotations is depicted.

**Figure 4.**
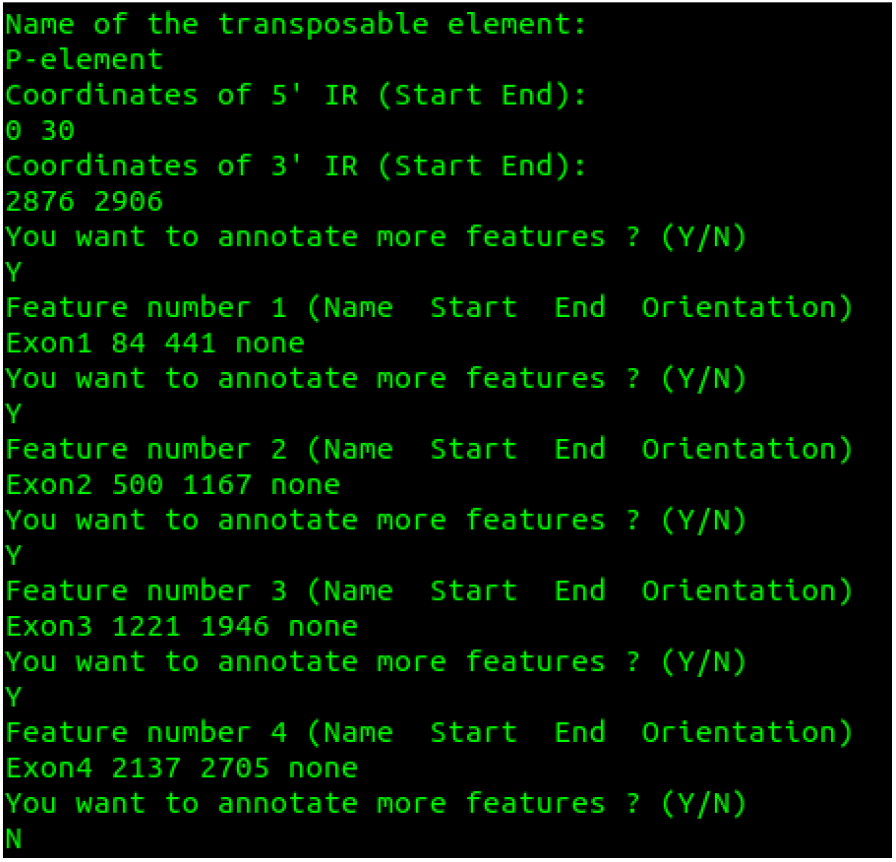
The Ubuntu terminal window showing the effect of *TE_annotator.sh* script when annotating the IR and exon regions of P-element.

**Figure 5.**
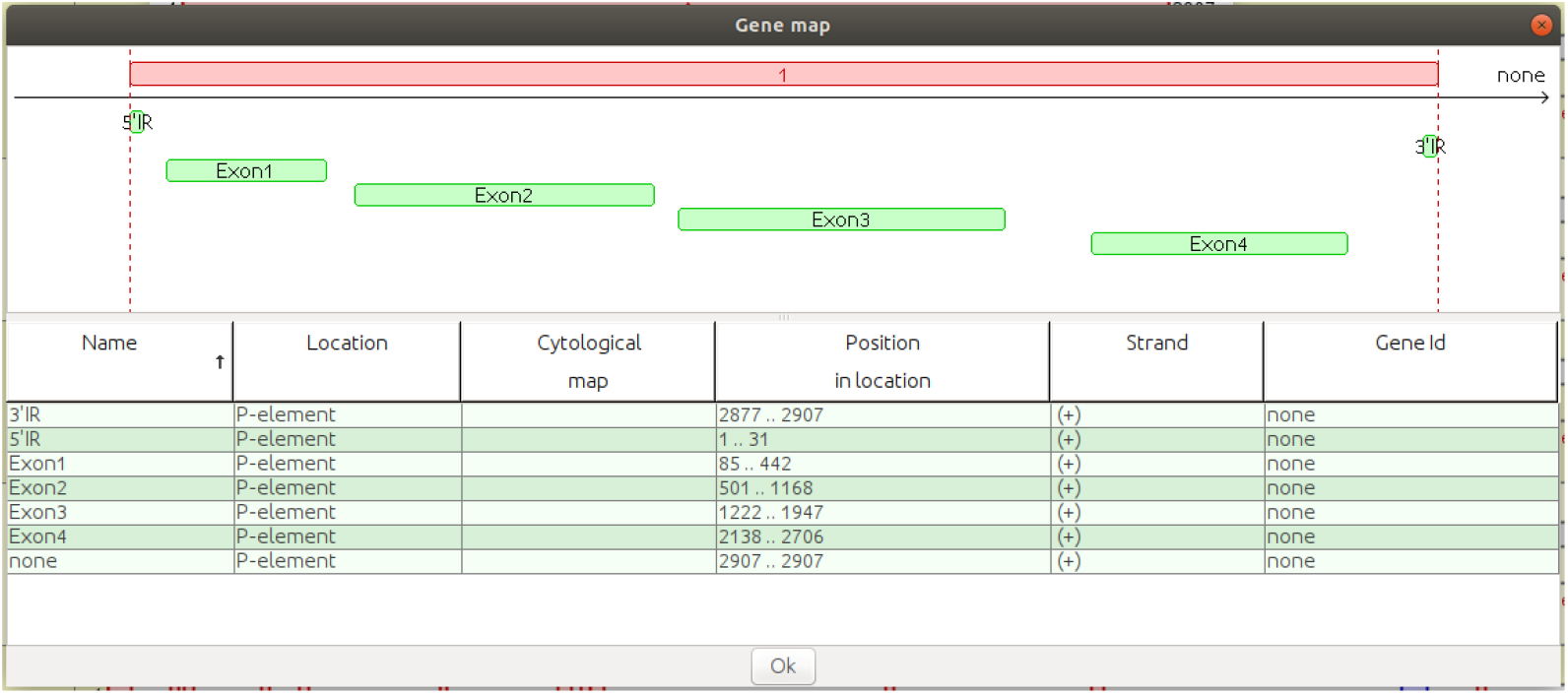
GA_v2 visualization of the annotation features implemented for P-element.

As previously stated (Ecovoiu *et al*., 2016), the annotation line referring to the first nucleotide of the sequence must start with the zero coordinate. Note that when starting from zero, each coordinate must be subtracted by one position.

All the annotations dealt by the two scripts are graphically depicted in green when they are overlapping with the query sequences.

### II. Case study

GA_v2 proved its efficiency when loaded with hundreds of trimmed Nanopore sequencing reads obtained from the genome of Horezu population of *D. melanogaster* (BioProject/NCBI, accession PRJNA629549) and containing P-element insertions.

Initially, we used the Ubuntu bash offline environment to perform a BLAST (Altschul *et al.,* 1990) alignment of the P-element reference sequence against the complete set of Nanopore reads. Consecutively, 212 reads containing P-element fragments were selected and fed to Geneious Prime 2020.2.4 (https://www.geneious.com). The 31 bp inverted repeat (IR) of the P-element was then searched against the 212 reads; 139 reads were reported by Geneious Prime as containing between 0 and 5 mismatches located in each IR. Out of them, 126 reads contained either 5’ or 3’ IRs of the P-element. For each of them we retrieved the respective IR plus the outflanking 200 nucleotides with Geneious Prime 2020.2.4 and the respective subsequences were fed to GA_v2 loaded with both r6.36 *D. melanogaster* release and P-element reference sequence.

GA_v2 detected 125 insertion coordinates and 19 out of them are reported at least twice, so mapped 106 unique TGNs. The export aggregate data table is presented in the Additional figure 1.

## Conclusions

Various tools are employed for TE annotations either in sequencing reads or in assembled genomes. To our knowledge, no such software was proved to be a perfect one, so there is a permanent bioinformatics work aimed to develop new tools or to optimize the integration of some of them in annotation pipelines (Ou *et al.,* 2019).

We report GA_v2 as reliable support tool for class II transposons annotation. GA_v2 kept all of the GA_1.19 performances, therefore the new version is also able to map TGN with the highest precision reported so far in literature. The ability to map self-insertions is also a unique asset of our tool (Ecovoiu *et al*., 2016) and is useful when detecting nested insertions and autoinsertions.

Herein we report the new features of GA_v2 designed to facilitate the high-throughput analysis of junction sequences data, regardless the sequences were generated by insertional mutation projects (mapping of artificial transposons) or by genome sequencing (mapping of natural TEs). The results obtained in a pilot experiment performed on Nanopore sequencing reads form *D. melanogaster* strain Horezu, Romania, point to the fact that GA_v2 was really useful when the class II transposon P-element was annotated.

The precision of GA_v2, along with its ability to map self-insertions and to generate consensus sequences for sets of TSDs or other flanking sequences recommend it to be considered along other TE annotation tools when annotation benchmarking test are run.

TE annotation benchmarking is a continuous challenge for TE community (Hoen *et al*., 2015). In this context, we consider that one aspect of this demanding endeavor would be to consider even the offline tools such as GA_v2 for comparative tests of performances, either when standing alone or when incorporated in pipelines dedicated to/for TE annotation.

---

**Additional figure 1.**
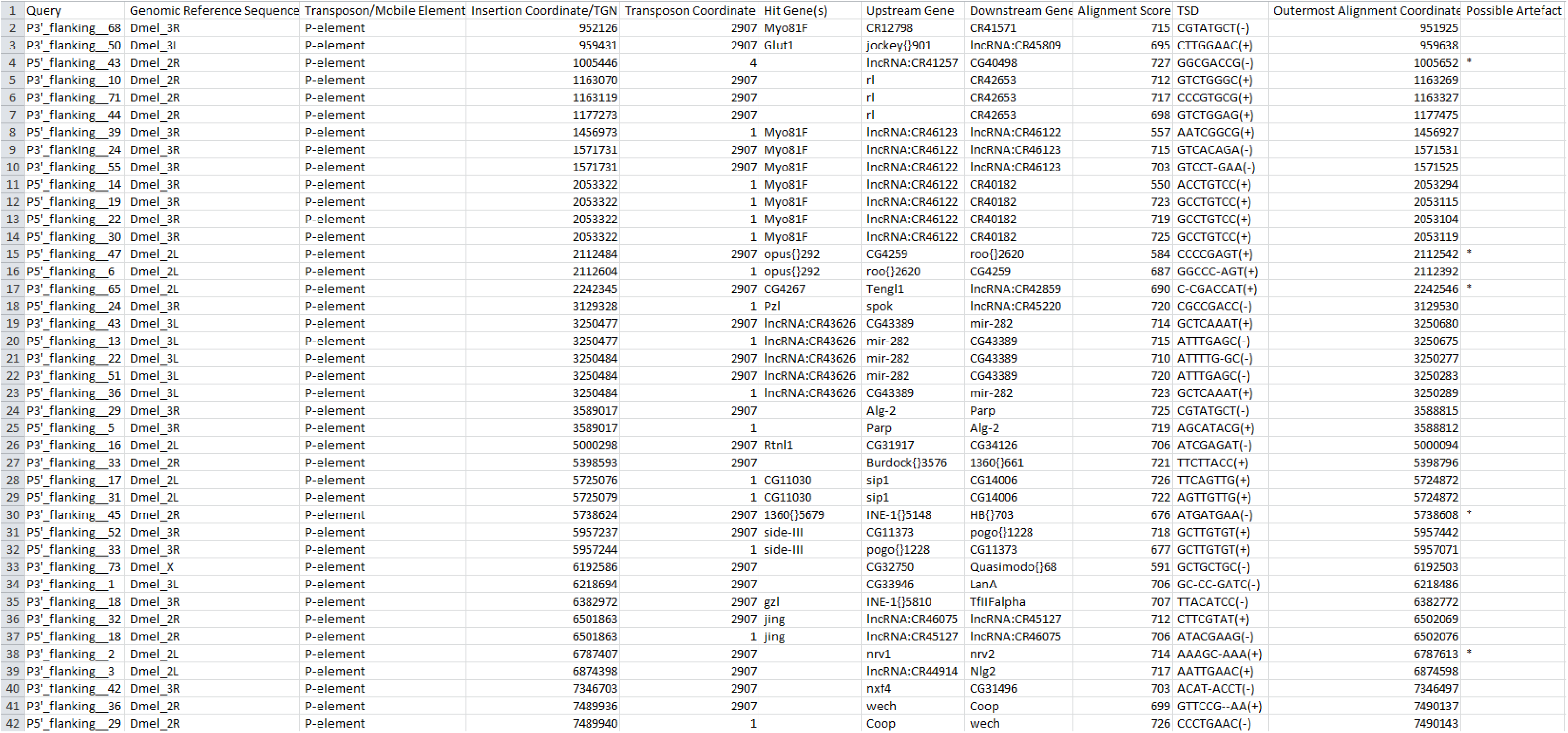
The tabular export of various genomic features identified by GA_v2 when mapping P-element in the genome of Horezu strain.

## References

1. Altschul, S. F., Gish, W., Miller, W., Myers, E. W., Lipman, D. J. (1990). Basic local alignment search tool. Journal of molecular biology, 215(3), 403–410. https://doi.org/10.1016/S0022-2836(05)80360-2

2. Bergman, C. M., Quesneville, H. (2007). Discovering and detecting transposable elements in genome sequences. Briefings in bioinformatics, 8(6), 382–392. https://doi.org/10.1093/bib/bbm048

3. Crooks, G. E., Hon, G., Chandonia, J. M., Brenner, S. E. (2004). WebLogo: a sequence logo generator. Genome research, 14(6), 1188–1190. https://doi.org/10.1101/gr.849004

4. Ecovoiu, A., Ghionoiu, I., Ciuca, A., Ratiu, A. C. (2016). Genome ARTIST: A robust, high-accuracy aligner tool for mapping transposon insertions and self-insertions. Mobile DNA, 7. https://doi.org/10.1186/s13100-016-0061-0

5. Geneious Prime 2020.2.4 (https://www.geneious.com)

6. Hoen, D. R., Hickey, G., Bourque, G., Casacuberta, J., Cordaux, R., Feschotte, C., Fiston-Lavier, A. S., Hua-Van, A., Hubley, R., Kapusta, A., Lerat, E., Maumus, F., Pollock, D. D., Quesneville, H., Smit, A., Wheeler, T. J., Bureau, T. E., Blanchette, M. (2015). A call for benchmarking transposable element annotation methods. Mobile DNA, 6, 13. https://doi.org/10.1186/s13100-015-0044-6

7. Linheiro, R. S., Bergman, C. M. (2012). Whole genome resequencing reveals natural target site preferences of transposable elements in *Drosophila melanogaster*. PloS one, 7(2), e30008. https://doi.org/10.1371/journal.pone.0030008

8. Liao, G., Rehm, E., Rubin, Gerald. (2000). Insertion site preferences of the P transposable element in *Drosophila melanogaster*. Proceedings of the National Academy of Sciences of the United States of America. 97. 3347–51. 10.1073/pnas.050017397. https://doi.org/10.1128/microbiolspec.MDNA3-0004-2014

9. Ou, S., Su, W., Liao, Y., Chougule, K., Agda, J., Hellinga, A. J., Lugo, C., Elliott, T. A., Ware, D., Peterson, T., Jiang, N., Hirsch, C. N., Hufford, M. B. (2019). Benchmarking transposable element annotation methods for creation of a streamlined, comprehensive pipeline. Genome biology, 20(1), 275. https://doi.org/10.1186/s13059-019-1905-y

10. Thurmond, J, Goodman, J. L., Strelets, V. B., Attrill, H., Gramates, L. S., Marygold, S. J., Matthews, B. B., Millburn, G., Antonazzo, G., Trovisco,V., Kaufman, T.C., Calvi, B.R., the FlyBase Consortium. (2019) FlyBase 2.0: the next generation. Nucleic Acids Research, 47(1), 759–765. https://doi.org/10.1093/nar/gky1003

11. Vendrell-Mir, P., Barteri, F., Merenciano, M., Gonzalez, J., Casacuberta, J. M., Castanera, R. (2019). A benchmark of transposon insertion detection tools using real data. Mobile DNA 10, 53. https://doi.org/10.1186/s13100-019-0197-9

12. www.ncbi.nlm.nih.gov/bioproject/?term=prjna629549

